# Comparison of transcriptional activation by corticosteroids of human MR (Ile-180) and haplotype (Val-180)

**DOI:** 10.1101/2024.12.08.627066

**Authors:** Yoshinao Katsu, Jiawen Zhang, Ya Ao, Patricia Donnellan, Michael E. Baker

## Abstract

While the human mineralocorticoid receptor (MR) regulates electrolyte homeostasis through aldosterone activation of the kidney MR, the MR also is highly expressed in the brain, where the MR is activated by cortisol. Here, we report the half-maximal response (EC50) and fold-activation by cortisol, aldosterone and other corticosteroids of human MR rs5522, a haplotype containing valine at codon 180 instead of isoleucine found in wild-type MR (Ile-180). MR rs5522 (Val-180) has been studied for actions in the human brain involving coping with stress and depression. We compared the EC50 and fold-activation by corticosteroids of MR rs5522 and wild-type MR transfected into HEK293 cells with either the TAT3 promoter or the MMTV promoter. Parallel studies investigated the binding of MR antagonists, spironolactone and progesterone, to MR rs5522 to investigate their use as antagonists of MR rs5522. In HEK293 cells with the MMTV promoter, MR rs5522 had a slightly higher EC50 compared to wild-type MR and a similar fold-activation for all corticosteroids. In contrast, in HEK293 cells with the TAT3 promoter, MR rs5522 had a higher EC50 (lower affinity) and fold-activation for cortisol compared to wild-type MR (Ile-180), while compared to wild-type MR, the EC50s of MR rs5522 for aldosterone and corticosterone were slightly lower and fold-activation was higher. Spironolactone and progesterone had similar antagonist activity for MR rs5522 and MR (Ile-180) in the presence of MMTV and TAT3 promoters in HEK293 cells indicating these antagonists are potential regulators of brain MR rs5522 to treat hyperactivity that contributes to Myalgic Encephalomyelitis/Chronic Fatigue Syndrome (ME/CFS).

## Introduction

The mineralocorticoid receptor (MR) belongs to the nuclear receptor family, which also contains other vertebrate steroid receptors: the glucocorticoid receptor (GR), progesterone receptor (PR), androgen receptor (AR) and estrogen receptor (ER) [1–4]. In humans and other terrestrial vertebrates, the MR functions to maintain electrolyte balance by regulating sodium and potassium transport in epithelial cells in the kidney and colon [5–8]. However, the MR also has important physiological functions in the brain, heart, skin and lungs [9–18].

Analysis of the human MR sequence by Arriza et al [1] revealed that, like other steroid receptors, the human MR is composed of four functional domains: a large amino-terminal domain (NTD) of about 600 amino acids, followed in the center by a DNA-binding domain (DBD) of about 65 amino acids, followed by a small hinge domain of about 60 amino acids that is connected to the ligand-binding domain of about 250 amino acids at the C-terminus, where corticosteroids, including aldosterone and cortisol bind to activate transcription [1,2,19–21].

Arriza et al. [1] also reported that the MR sequence was related to the glucocorticoid receptor (GR) sequence, consistent with evidence that some corticosteroids, such as cortisol and corticosterone were transcriptional activators of both the MR and GR [9,11,15,20,22] and that aldosterone, cortisol, corticosterone and 11-deoxycorticosterone have similar binding affinity for human MR [1,15,22–24]. Activation by cortisol and corticosterone of the MR by two steroids that are ligands for the GR [24–26], is consistent with the evolution of the GR and MR from a common ancestral corticoid receptor (CR) in a cyclostome, a jawless fish that evolved about 550 million years ago at the base of the vertebrate line [20,27–29].

Humans contain three almost identical MR transcripts that differ at codons 180 and 241 in the NTD [30] (Fig 1). Human MR (Ile-180), cloned by Arriza et al [1], has been extensively studied for transcriptional activation by aldosterone and other corticosteroids [1,22,23,25,26,30]. Here, we focus on the response to corticosteroids of another human MR, haplotype rs5522 (Val-180), which contains a valine at codon 180 [31–34] (Figure 1). rs5522 (Val-180) mediates the physiological responses of steroids on stress and depression in humans [33–37] making transcriptional activation by steroids of rs5522 of much interest.

**Fig 1.**
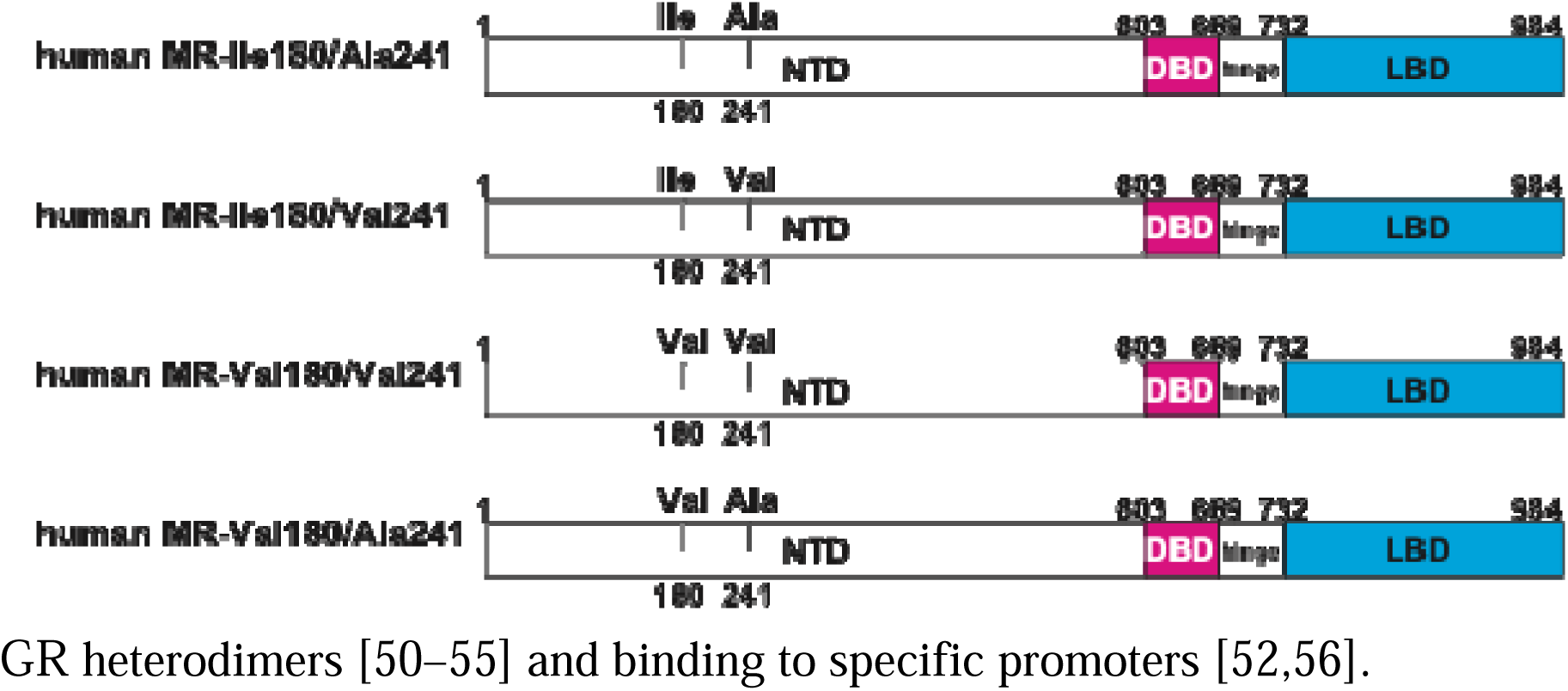
Human MR Genes. There are three human MR genes with 984 amino acids in GenBank. These MRs contain either isoleucine-180, alanine-241 (Ile-180/Ala-241: accession AAA59571), isoleucine-180, valine-241 (Ile-180/Val-241: accession XP_054206038), or valine-180, valine-241 (Val-180/Val-241: accession NP_000892), in the NTD. The fourth MR isoform is rs5522, which contains valine-180 instead of isoleucine-180 (Val-180/-Ala241) [31–33]. Isoleucine-180, alanine-241 (Ile-180/Ala-241: accession AAA59571) is the classical MR that regulates electrolyte transport in the kidney. The MR also functions in the brain. A mutant MR, rs5522, which contains valine-180 instead of isoleucine-180 (Val-180/-Ala241) [31–33] is found in a patient with chronic fatigue syndrome (ME/CFS).

Aldosterone [32,33] and cortisol [33,38] are transcriptional activators of MR rs5522. However, EC50s and fold-activation by aldosterone, corticosterone and 11-deoxycorticosterone for the rs5522 haplotype have not been reported, which is our goal. Activation of rs5522 by a corticosteroid other than cortisol could provide clues to the unique physiological actions mediated by rs5522 in the brain MR. Nor has the binding of spironolactone and progesterone, two MR antagonists [39,40] to rs5522 been investigated. Inhibition of brain rs5522 would provide a selective method for treating hyperactivity mediated by brain rs5522, which contributes to Myalgic Encephalomyelitis/Chronic Fatigue Syndrome (ME/CFS). To provide this information, we investigated the effect of corticosteroids and of mineralocorticoid antagonists on transcriptional activation of rs5522 in presence of two promoters: TAT3 [41,42] and MMTV [19,43]. We find that in HEK293 cells with the MMTV promoter, each corticosteroid had a similar EC50 and fold-activation for rs5522 and wild-type MR (MR Ile-180). In contrast, in HEK293 cells with the TAT3 promoter, compared to wild-type MR, rs5522 had a slightly lower EC50 for aldosterone and corticosterone and a higher EC50 for cortisol and a similar EC50 for 11-deoxycorticosterone. However, fold-activation for cortisol activation of rs5522 was higher than for MR (Ile-180) in the presence of TAT3, which is interesting because cortisol is the biological ligand for the brain MR [13,15,17,18,44]. We find that spironolactone and progesterone had similar antagonist activity for rs5522 and MR (Ile-180) in the presence of the MMTV and TAT3 promoters in HEK293 cells, this supports investigating the use of spironolactone for treating patients with ME/CFS pathologies due to rs5522. The absence of a large difference in EC50 and/or fold-activation between rs5522 and (MR Ile-180) in their response to corticosteroids suggests that other mechanisms are important in rs5522 regulation of the response to stress and depression. These may include post-translational modification of the NTD [45–49], formation of MR-GR heterodimers [50–55] and binding to specific promoters [52,56].

## Results

### Corticosteroid-dependent activation of wild-type human MR (Ile-180) and rs5522 (MR Val-180)

We have used human MR (accession AAA59571) [1], which contains isoleucine at codon 180 for our studies of the human MR [23,30]. For this project, as described in the Methods Section, we constructed rs5522 from human MR (Ile-180, Ala-241). In Fig 2, we show the concentration dependence of transcriptional activation by corticosteroids of rs5522 and MR (Ile-180) transfected into HEK293 cells with either an MMTV or a TAT3 luciferase promoter. Luciferase levels were used to calculate an EC50 and fold-activation for each steroid (Table 1).

**Fig 2.**
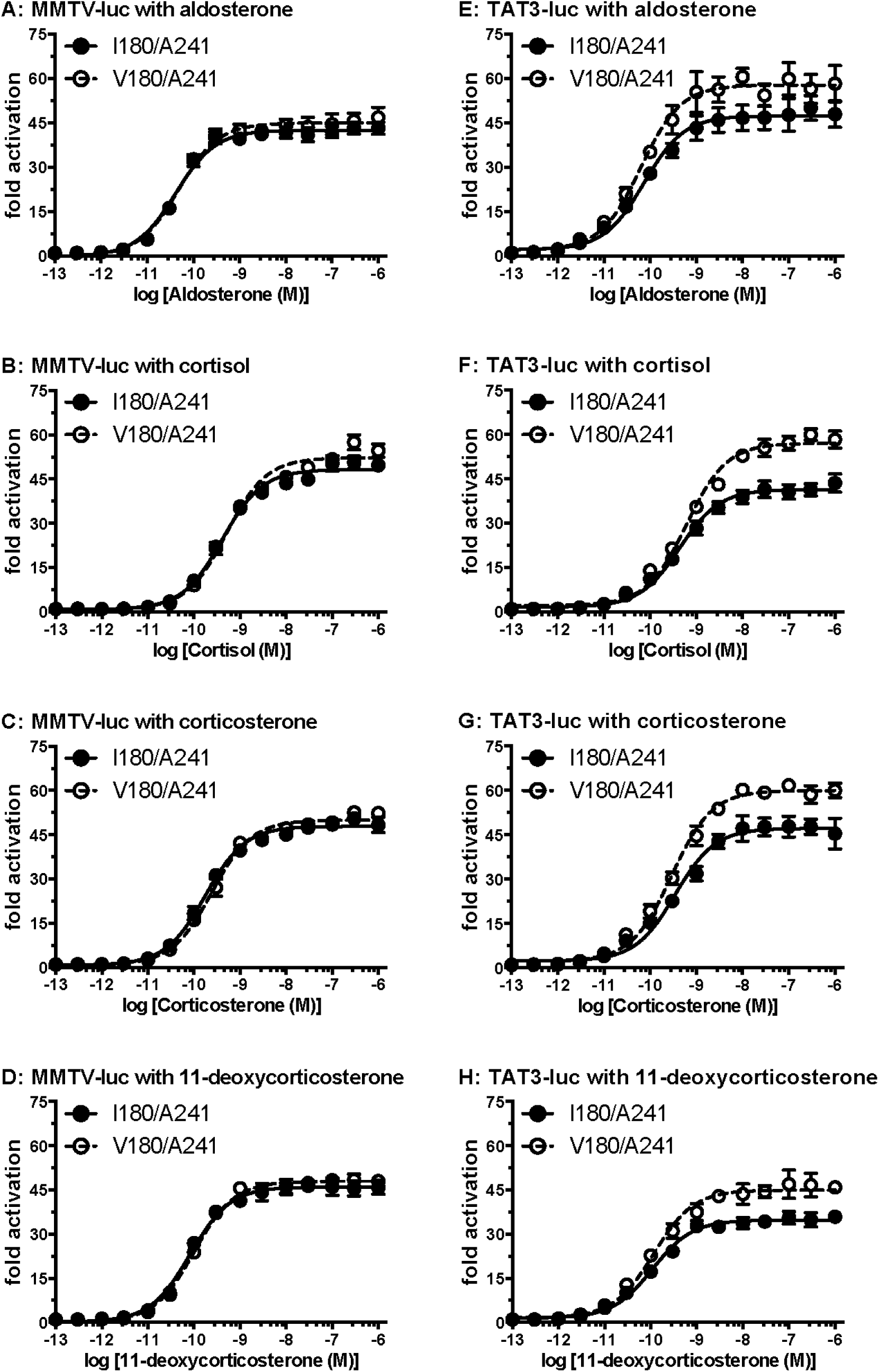
Concentration-dependent transcriptional activation by corticosteroids of wild-type human MR (Ile-180) and rs5522 (MR Val-180) in the presence of either the MMTV promoter or the TAT3 promoter. Plasmids for wild-type MR (Ile-180) and rs5522 (MR Val-180) were expressed in HEK293 cells with either an MMTV-luciferase promoter (2A-2D) [19,43] or a TAT3-luciferase promoter (2E-2H) [2,41,42]. Cells were treated with increasing concentrations of either aldosterone, cortisol, 11-deoxycorticosterone, corticosterone, or vehicle alone (DMSO). Results are expressed as means ± SEM, n=3. Y-axis indicates fold-activation compared to the activity of vector with vehicle (DMSO) alone as 1. A. MR (Ile-180) and rs5522 (MR Val-180) with aldosterone. B. MR (Ile-180) and rs5522 (MR Val-180) with cortisol. C. MR (Ile-180) and rs5522 (MR Val-180) with corticosterone. D. MR (Ile-180) and rs5522 (MR Val-180) with 11-deoxycorticosterone. E. MR (Ile-180) and rs5522 (MR Val-180) with aldosterone. F. MR (Ile-180) and rs5522 (MR Val-180) with cortisol. G. MR (Ile-180) and rs5522 (MR Val-180) with corticosterone. H. MR (Ile-180) and rs5522 (MR Val-180) with 11-deoxycorticosterone.

**Table 1.** Steroid Activation of Human MRs in HEK293 Cells with an MMTV or a TAT3 Promoter.

|  |  | MMTV-luc |  | TAT3-luc |  |
| --- | --- | --- | --- | --- | --- |
|  |  | **Wild-type | **MR-rs5522 | **Wild-type | **MR-rs5522 |
| Aldosterone | EC50 (nM) | 0.042 | 0.048 | 0.074 | 0.063 |
|  | *Fold Activation (± SEM) | 41.9 (± 1.9) | 44.5 (± 3.5) | 47.7 (± 5.6) | 59.9 (± 5.4) |
|  | ***Ratio values | 1 | 1.06 | 1 | 1.26 |
| Cortisol | EC50 (nM) | 0.41 | 0.51 | 0.43 | 0.62 |
|  | *Fold Activation (± SEM) | 50.3 (± 2.6) | 51.7 (± 0.9) | 40.6 (± 2.4) | 57.1 (± 2.4) |
|  | ***Ratio values | 1 | 1.03 | 1 | 1.41 |
| Corticosterone | EC50 (nM) | 0.18 | 0.24 | 0.35 | 0.29 |
|  | *Fold Activation (± SEM) | 48.5 (± 1.7) | 49.2 (± 0.9) | 47.8 (± 3.5) | 61.7 (± 0.06) |
|  | ***Ratio values | 1 | 1.01 | 1 | 1.29 |
| 11-deoxy-corticosterone | EC50 (nM) | 0.083 | 0.1 | 0.11 | 0.12 |
|  | *Fold Activation (± SEM) | 46.1 (± 3.0) | 48.3 (± 1.2) | 35.6 (± 2.1) | 47.0 (± 4.8) |
|  | ***Ratio values | 1 | 1.05 | 1 | 1.32 |

\*Fold Activation values were calculated based on the 100 nM values.
\*\* MR-rs5522= V180/A241, Wild-type MR = Ile180, A241
\*\*\*Ratio values were calculated by dividing the fold-activation value of MR-rs5522
(V180/A241) form by the value of Wild-type (I180/A241) form.

Fig 2 reveals that in HEK293 cells, transfected with the MMTV promoter, the EC50 and fold-activation values were similar for aldosterone, cortisol, corticosterone and 11-deoxycorticosterone. It is in the experiments with TAT3 promoter in HEK293 cells that we find differences between rs5522 and MR (Ile-180).

As shown in Fig 2 and Table 1, in HEK293 cells transfected with TAT3, and with either aldosterone or corticosterone had a slight decrease in EC50 (stronger binding) for rs5522 compared to MR (Ile-180), while cortisol had a slight increase in EC50 for rs5522 compared to MR (Ile-180). The EC50 of 11-deoxycorticosterone was about the same value for rs5522 and MR (Ile-180). Also as shown in Fig 2 and Table 1, in HEK293 cells transfected with TAT3, aldosterone increased fold-activation by about 25% for rs5522 compared to MR (Ile-180). Cortisol, which activates the MR in organs that lack 11β-hydroxysteroid dehydrogenase-type 2 [5,57–59], increases fold activation by about 40% for rs5522 compared to MR (Ile-180). Corticosterone and 11-deoxycorticosterone increased fold-activation by about 30% for rs5522 compared to MR (Ile-180).

### Spironolactone and progesterone are antagonists for rs5522 and MR (Ile-180)

Spironolactone and progesterone are widely used MR (Ile-180) antagonists [39,49]. To determine if spironolactone and progesterone are antagonists for rs5522, we added either spironolactone or progesterone to HEK293 cells transfected with either human MR (Ile-180) or rs5522 and either an MMTV promoter or a TAT3 promoter, which were then incubated with 0.1 nM aldosterone. Fig 3 shows that 10 nM spironolactone or 10 nM progesterone inhibit activation of rs5522 and MR (Ile-180) by 0.1 nM aldosterone.

**Fig 3.**
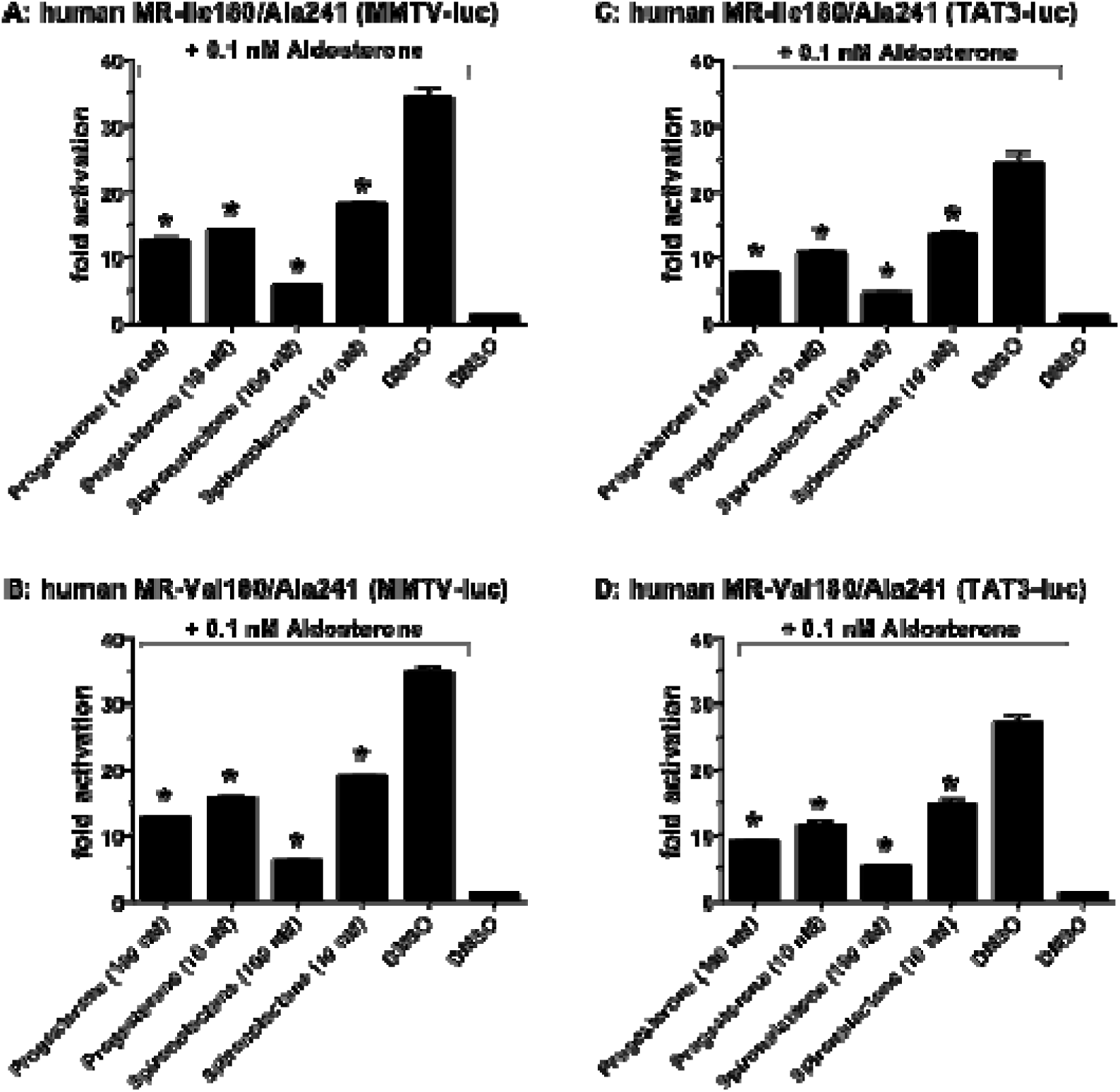
Inhibition of Aldosterone Activation rs5522 and human MR (Ile-180, Ala-241) by spironolactone and progesterone. HEK293 cells were transfected with either human MR (Ile-180, Ala-241) or rs5522 (Val-180, Ala-241) and either the MMTV promoter or the TAT3 promoter. Then these cells were incubated with either 10 nM or 100 nM spironolactone or progesterone for 1 hour and then 0.1 nM aldosterone was added and the incubation continued for 24 hrs. HEK293 cells were harvested and processed for luciferase. Results are expressed as means ± SEM, n=3. Y-axis indicates fold-activation compared to the activity of vector with vehicle (DMSO) alone as 1.

### FLAG-tagged expression of human MR (Ile-180) and rs5522

To investigate the protein expression levels of FLAG-tagged MR (Ile-180) and rs5522 (Val-180), HEK293 cells transfected with each construct were collected and treated with SDS sample buffer. Subsequently, FLAG-tagged proteins were subjected to SDS-PAGE on a 10% polyacrylamide gel and transferred to an Immobilon membrane and detected using an antibody against the FLAG tag. We detected a single band of human MR (Ile-180) and rs5522 (Fig 4), which were used in the experiments depicted in Fig 2. This analysis of FLAG-tagged human MR (Ile-180) and rs5522 (Val-180) confirms that there were similar protein expression levels of these two MRs in our experiments.

**Fig 4.**
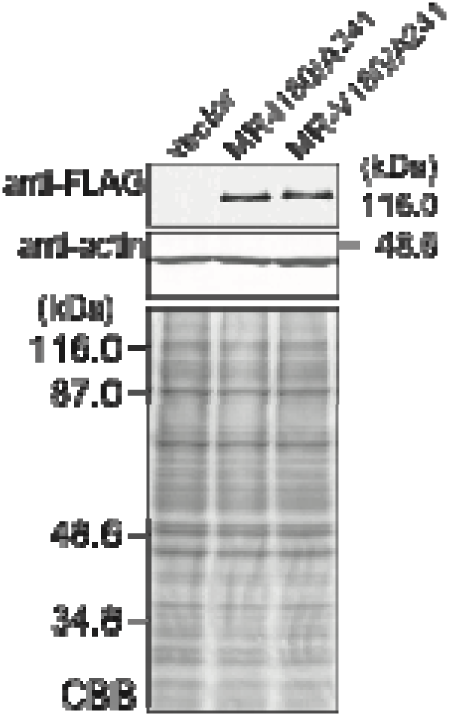
FLAG-tagged expression of human MR (Ile-180) and rs5522 (Val-180). HEK293 cell lysates transfected with FLAG-tagged MR (Ile-180, Ala-241) and rs5522 (Val-180, Ala-241) were treated with sample buffer and applied to a 10% SDS-polyacrylamide gel and then transferred onto a membrane. The expressed FLAG-tagged proteins were detected with anti-FLAG antibody. The expression level of endogenous actin was measured using an actin antibody. Upper panel: Western blot. Lower panel: Coomassie blue stain (CBB).

## Discussion

The cloning of the human MR by Arriza et al [1] provided tools for investigating the function of the MR in diverse tissues other than the distal tubule of the kidney, including the MR in the hippocampus, which is receiving increasing attention as an MR-responsive organ [14,15,17,18]. Indeed, the MR in the hippocampus is downregulated by extended stress and depression [15,35,60,61], and MR synthesis is induced by anti-depressants. Increased activity of the MR inhibits activity of the hypothalamic-pituitary-adrenal axis and promotes slow wave cognition, reducing anxiety.

The MR rs5522 (Val-180), which is the focus of this report, is intriguing and important because humans with MR (Val-180) have increased response to stress [17,18,33,36] despite the presence of only a single amino acid difference between the rs5522, which contains valine at codon 180, instead of isoleucine at codon 180 as found in wild-type MR [17,35,36,62]. Such a profound physiological effect of a point mutation in the NTD, which is distant from the LBD and DBD is unexpected. Moreover, valine-180 is not located in one of the transcriptional activation domains in the NTD, which occur at residues 1-169 and 451-603 in the NTD of human MR [63,64].

As a first step towards understanding the biological actions of the Ile180Val mutation in human MR, we investigated transcriptional activation of MR (Val-180) by a panel of corticosteroids including aldosterone and cortisol, for comparison with activation of wild-type human MR, in the presence of either the MMTV promoter or the TAT3 promoter [Figure 2, Table 1]. Interestingly, in cells with TAT3, but not in cells with MMTV, there were differences in transcriptional activation of MR (Val-180) and wild-type MR (Ile-180) by corticosteroids.

Among the corticosteroids, cortisol is of interest because in the human brain cortisol is the physiological steroid for the MR because the circulating cortisol concentration is about 200-fold higher compared to aldosterone, and11β-hydroxysteroid dehydrogenase-type 2, which can inactivate cortisol [57–59], is absent in the hippocampus [57]. Cortisol had a higher EC50 for MR (Val-180) than for wild-type MR in HEK293 cells transfected with TAT3, in agreement with [33]. Interestingly, fold-activation of MR (Val-180) by cortisol was about 40% higher than for cortisol activation of wild-type MR. Aldosterone had a slightly lower EC50 and 26% higher fold-activation for MR (Val-180) compared to wild type MR. Corticosterone and 11-deoxycorticosterone had slightly lower EC50s and about 30% higher fold-activation for MR (Val-180) than for wild-type MR.

We also found that spironolactone and progesterone, which inhibit wild-type MR also inhibited aldosterone activation of MR (Val-180) [Figure 3]. This inhibition of MR (Val-180) provides an explanation for understanding the basis for a recent clinical case in which treatment with spironolactone cured a patient with hyperactivity and chronic fatigue syndrome (ME/CFS). Other mineralocorticoid receptor antagonists may also be useful in treating MR (Val-180)-dependent hyperactivity.

The role of the weaker EC50 and higher fold-activation of cortisol for MR (Val-180) by themselves may be insufficient to support the profound biological response to stress and depression in people with MR (Val-180) [15,35,36,60,61]. We intend to investigate other mechanism(s) that may contribute to novel physiological activity of MR (Val-180). One possibility arises from the different response to cortisol between MR (Val-180) and wild-type MR in the presence of either TAT3 or MMTV [30,65], and the similar finding by DeRijk et al. [33] in the response to cortisol of MR (Val-180) in cells with either TAT3 or MMTV promoters. It may be that other promoters in the brain exert larger differences between MR (Val-180) and wild-type MR in transcriptional activation by cortisol and other corticosteroids, which may be physiologically important.

Another possible mechanism is regulation of cortisol-mediated transcription of MR (Val-180) by phosphorylation of one or more serine residues near codon 180 the NTD on the MR [46–49,66]. Indeed, Shibata et al [48,66], Kino et al. [49] and Faresse et al. [67] have identified several phosphorylation sites in the NTD of human MR. These sites include Ser-129, Ser-183, Ser-196, Ser-227, Ser238, Ser-250 and Ser-263. Substitution of one or more serines in MR (Val-180) with alanine or aspartic acid would determine whether phosphorylation is important in corticosteroid activation of MR (Val-180).

We also will investigate the effect of the formation of heterodimers between MR (Val-180) and the GR [50–55,68] on activation of MR (Val-180), which is relevant to MR activity in the brain based on research by Mifsud and Ruel [52], who found that stress promotes MR-GR heterodimers in the hippocampus. Promotion by stress of heterodimers between MR (Val-180) and the GR may be important in the effects of MR (Val-180) on stress and depression.

## Materials & Methods

### Construction of plasmid vectors

Full-length mineralocorticoid receptor (MR) of human as registered in Genbank (accession number: XP_054206038) was used as wild-type human MR. MR rs5522 was constructed by the replacement of isoleucine-180 by valine using KOD-Plus-mutagenesis kit (TOYOBO). The nucleic acid sequences of all constructs were verified by sequencing.

### Chemical reagents

Cortisol, corticosterone, 11-deoxycorticosterone, aldosterone, progesterone and spironolactone were purchased from Sigma-Aldrich. For reporter gene assays, all hormones were dissolved in dimethyl-sulfoxide (DMSO); the final DMSO concentration in the culture medium did not exceed 0.1%.

### Transactivation assays and statistical analyses

Transfection and reporter assays were carried out in HEK293 cells, as described previously [23]. HEK293 cells were seeded in 24-well plates at 5×104 cells/well in phenol-red free Dulbecco’s modified Eagle’s medium supplemented with 10% charcoal-stripped fetal calf serum (HyClone). After 24 hours, the cells were transfected with 100 ng of receptor gene, reporter gene containing the *Photinus pyralis* luciferase gene and pRL-tk, as an internal control to normalize for variation in transfection efficiency; pRL-tk contains the *Renilla reniformis* luciferase gene with the herpes simplex virus thymidine kinase promoter. Each assay had a similar number of cells, and assays were done with the same batch of cells in each experiment. All experiments were performed in triplicate. Promoter activity was calculated as firefly (*P. pyralis*)-lucifease activity/sea pansy (*R. reniformis*)-lucifease activity. The values shown are mean ± SEM from three separate experiments, and dose-response data, which were used to calculate the half maximal response (EC50) for each steroid, were analyzed using GraphPad Prism.

### Antibodies and Western blotting of FLAG-tagged MR

Primary antibodies for Western blotting were purchased and used at the indicated dilutions: anti-FLAG (clone #1E6, FUJIFILM Wako Pure Chemical Corporation, 1:750), and anti-betaActin (clone # 6D1, Medical & Biological Laboratories Co., LTD., 1:3000). Cells transfected as described in “Transactivation assays and statistical analysis” were directly lysed with sodium dodecyl sulfate (SDS) sample buffer and subjected to SDS-PAGE on a 10% polyacrylamide gel and transferred to an Immobilon membrane. The membrane was blocked with 10% skim milk overnight at 4 °C and then incubated with a primary antibody for 4 hours at room temperature. The membrane was washed with TTBS (0.1% Tween-20 in Tris-buffered saline) for at three times for 10 min and incubated with AP-conjugated secondary antibody (COSMO BIO Co. Ltd.) for 1 hour at room temperature. After the final wash with TTBS, the membrane was treated with NBT-BCIP solution). The gels were also stained with Coomassie Blue.

## Contributions

Yoshinao Katsu: Investigation, Conceptualization, Supervision, Formal Analysis, Writing – original draft, Writing – review & editing.

Jiawen Zhang: Data curation, Investigation, Methodology. Ya Ao: Data curation, Methodology.

Patricia Donnellan: Conceptualization, Formal analysis, Writing – review & editing.

Michael E. Baker: Conceptualization; Formal analysis, Supervision, Writing – original draft, Writing – review & editing.

